# Genetic aberrations in chromatin factors are pervasive, mutually exclusive, associated with a high mutational burden and improved response to checkpoint immunotherapy in cancer

**DOI:** 10.1101/861823

**Authors:** Martina Markovska, Rodney Rosalia, Aleksandra Stajkovska, Sanja Mehandziska, Marija Janevska, Ivan Kungulovski, Zan Mitrev, Goran Kungulovski

**Author notes:** Corresponding author Dr. Goran Kungulovski, Bio Engineering LLC, Laboratory of Genetics and Personalized Medicine, Zan Mitrev Clinic, Ivan Agovski 7-1, 1000, Skopje, Republic of Macedonia.

## Abstract

In recent years, it has been shown that many of the pervasive genetic defects in cancer occur in chromatin modifiers. Herein, we analyzed the distribution and mutual relationships of genetic aberrations in relevant chromatin modifiers in tens of thousands of publicly available cancer datasets. We observed that in general, they are mutually exclusive, and their prevalence is higher in some types of cancers compared to others. Moreover, we observed a strong association between aberrations in selected chromatin modifiers and tumor mutational burden, leading to an improved response to checkpoint immunotherapy. All in all, this study uncovered interesting relationships between chromatin factors and features of cancer, which warrant follow up functional and clinical studies.

## BACKGROUND

Cancer denotes a multifaceted group of diseases, generally characterized by cell regulation failure, which results in abnormal and uncontrolled cell growth. These properties equip the cells with the potential to divide in an uncontrolled manner, hide from the immune system, and over time, invade and spread into neighboring tissues and organs of the body [1]. Cancer is thought to be driven by the accumulation of genetic defects such as mutations, deletions, amplifications and translocations in so-called oncogenes and tumor-suppressor genes [2]. Therefore, historically, most of the endeavors undertaken in cancer biology have been focused on genetic aberrations in genes with prominent roles in DNA repair, cell proliferation, and cell signaling. However, in the past years, with the broad application of NGS sequencing technologies for comprehensive cancer profiling it has been shown that many of the pervasive genetic defects in cancer occur in epigenetic (chromatin) regulators [3]. Such mutations in chromatin writers, readers, erasers, remodelers and even the histone proteins themselves, can lead to global alterations in the chromatin of cells, which in turn can steer the cells to altered epigenetic states and expression programs that contribute to tumorigenesis [4-6]. This indicates that genetic alterations of chromatin regulators provide the cancer cell with a very efficient means to rewire its transcriptional programs and adapt quickly to microenvironmental or therapeutic pressures [7]. In addition, simply because of the sheer ubiquity of mutations in epigenetic modifiers in most types of cancer, they are increasingly regarded as novel targets for cancer treatment. Hence, obtaining a greater understanding and deeper insights into the frequency, distribution, mutual relationships and features of these regulators in cancer can significantly aid our understanding of cancer biology. In this study, we sought to analyze the distribution, mutual relationships and features of selected 57 chromatin modifiers in tens of thousands of available genetic maps of different types of cancers.

## RESULTS

The presence of genomic alterations in chromatin modifiers is pervasive in cancer We tested the prevalence and distribution of genomic alterations in 57 genes selected from literature [8, 9] coding for chromatin modifiers in 158 non-redundant studies encompassing 31254 tumor samples from the TCGA study and others, as well as 28 genes in 10945 tumor samples from the MSK-IMPACT study [10] **(Additional File 1, Suppl. Figures 1 and 2)** deposited in the cBioportal platform [11, 12]. Our preliminary analysis revealed that chromatin remodelers were the most frequently altered epigenetic category of regulators, followed by histone-modifying, and DNA modifying enzymes **(Figure 1; Additional File 1, Suppl. Figures 3 and 4)**, with mutations (missense and truncating) being the most common genomic alterations **(Additional File 1, Suppl. Figure 3C)**. As a proof of principle, we have detected readily known aberrations such as enrichment of *PBRM1* in renal cancer, and enrichment of *ATRX* with *IDH1* in gliomas [13] **(Figure 1)**. Particular types of cancers (e.g lung cancer, bladder cancer, diffuse glioma, uterine cancer, skin cancer) had a higher incidence of genomically altered chromatin modifiers in contrast to others (e.g rhabdoid cancer, thyroid cancer, peripheral nervous system cancers, Wilms tumors) **(Figure 1; Additional File 1, Suppl. Figure 2)**.

**Figure 1.**
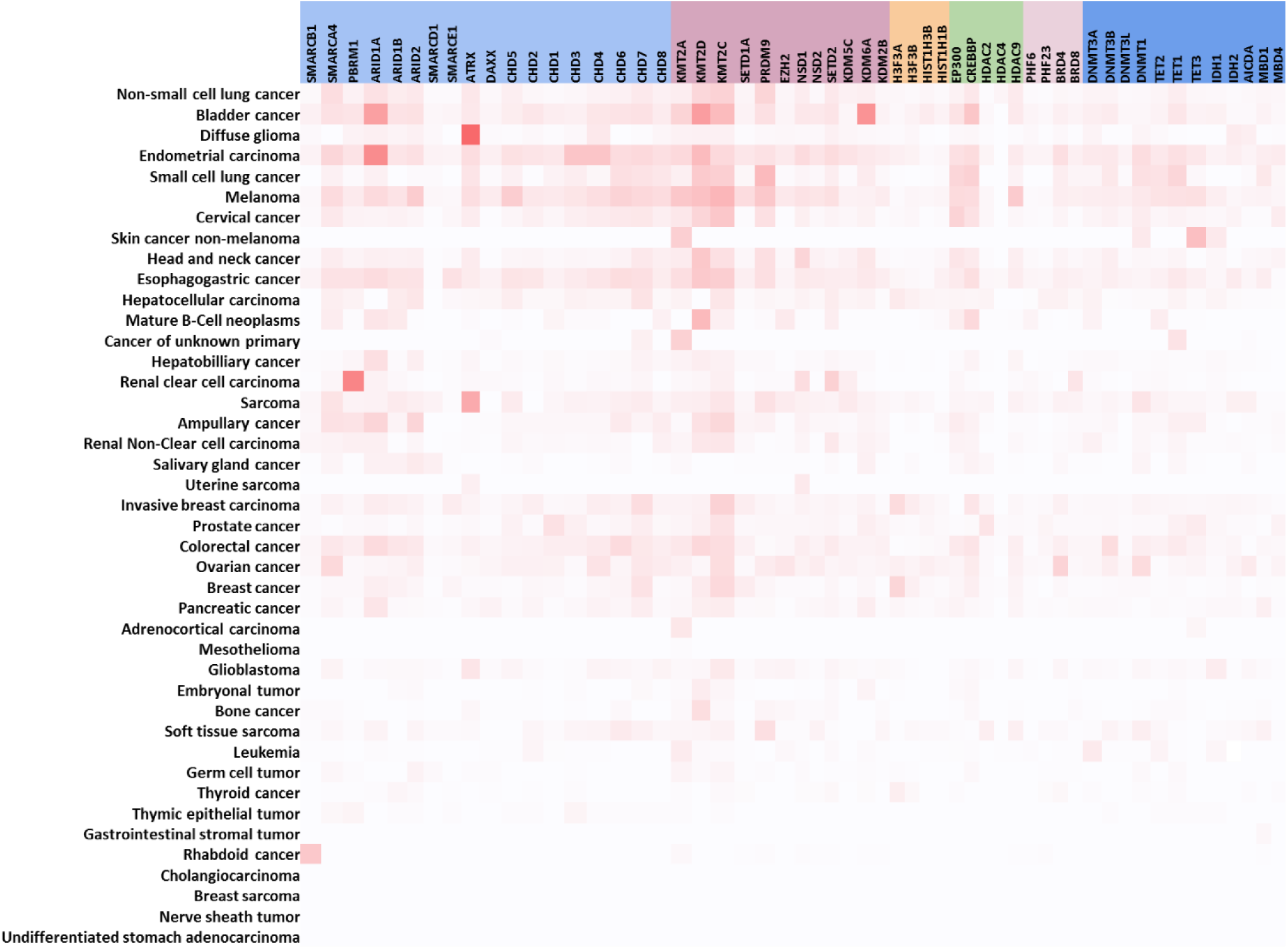
Prevalence and relationships between genomic alterations in chromatin modifiers; Heatmap showing the prevalence of alterations in chromatin modifiers per cancer type.

### Genomic alterations in chromatin modifiers are mutually exclusive in cancer

To get a better insight of the mutual relationships between genomically altered chromatin modifiers, we wanted to study their finer distribution in available cancer maps taken from patients and cancer cell lines **(Figure 2; Additional File 1, Suppl. Figures 5 and 6)**. The vast majority of altered chromatin modifiers followed a trend of mutual exclusivity in the TCGA datasets, as well as MSK-IMPACT and cancer cell lines as visualized by oncoprint analysis, and the lack of or very weak correlation **(Figure 2; Additional File 1, Suppl. Figures 7-9; Additional File 2)**. Contrary to the general trend of mutual exclusivity, we noted a couple of exceptions of co-occurrence that can be readily visualized by oncoprint, and were moderately correlated **(Figure 2; Additional File 1, Suppl. Figures 7-9; Additional File 2 and 4)**, for instance, *HISTH1B-HIST1H3B-DAXX* (Log2 Odds ratio >3, P-value <0.001); *CHD3-PHF23* (CNAs, Spearman ρ =0.77, Log2 Odds ratio >3, P-value <0.001), *EZH2-KMT2C* (CNAs, Spearman ρ =0.68, Log2 Odds ratio >3, P-value <0.001); *BRD4-DNMT1-SMARCA4* (Log2 Odds ratio >3, P-value <0.001); *PBRM1-SETD2* (CNAs, Spearman ρ =0.52, Log2 Odds ratio >3, P-value <0.001); *ATRX-IDH1* (Mutations, Spearman ρ =0.34, Log2 Odds ratio >3, P-value <0.001), as well as *KMT2C/KMT2D* with chromatin remodelers and histone acetyltransferases **(Additional File 1, Suppl. Figures 7-11; Additional File 4)**. These co-occurring alterations in chromatin modifiers were even more visible in selected cancer types **(Additional File 1, Suppl. Figures 12-16)**, such as deletion/amplifications of *SMARCA4* and *DNMT1* in lung cancer **(Additional File 1, Suppl. Figure 12)**, amplifications of *SMARCA4, DNMT1* and *BRD4* in gliomas/glioblastomas and uterine cancers **(Additional File 1, Suppl. Figures 14 and 16)**. In the same vein, we observed a pervasive overlap of alterations in histone-modifying genes such as *KMT2C, KMT2D, KDM6A, CREBBP, EP300* with chromatin remodelers and other chromatin modifiers in bladder cancer **(Additional File 1, Suppl. Figure 13)**, and deletions of *HDAC2* and *ARID2* in melanoma **(Additional File 1, Suppl. Figure 15)**.

**Figure 2.**
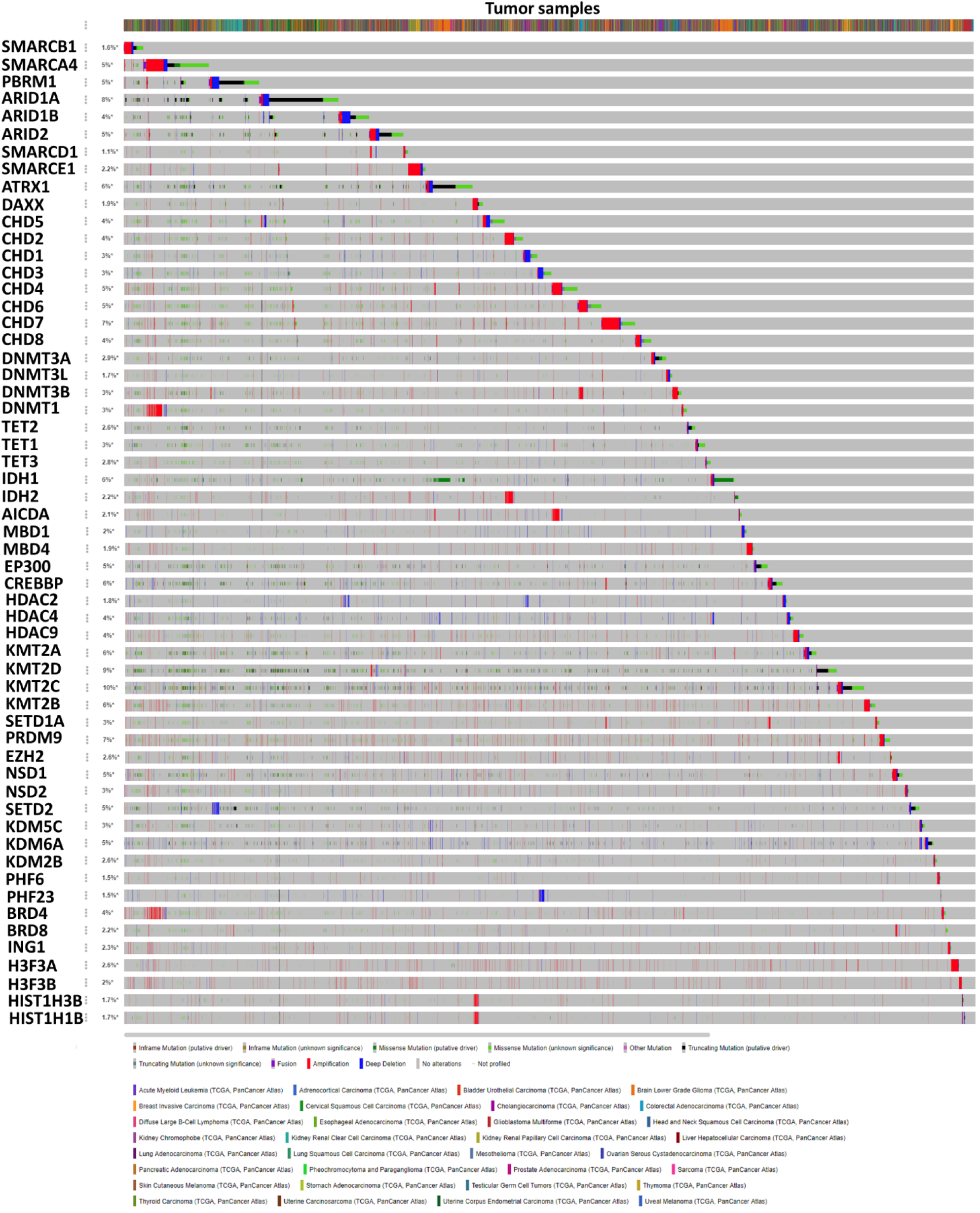
Prevalence and relationships between genomic alterations in chromatin modifiers; oncoprint analysis showing a general trend of mutual exclusivity between alterations of chromatin modifiers. Cancers lacking genomic alteration in chromatin modifiers are not shown.

### Genomic alterations in selected chromatin modifiers are associated with higher TMB and an improved response to checkpoint immunotherapy

Next, we sought to investigate the relationships between genomically altered chromatin modifiers and other molecular and clinical features, such as tumor mutational burden (TMB), and patient survival. Interestingly, cancers with high mutation rates **(Additional File 1, Suppl. Figure 17A and B)**, such as endometrial cancer, colorectal cancer, skin cancer, glioma, lung cancer, and bladder cancer were the types of cancers with a high prevalence of genomically altered chromatin modifiers **(Figure 1; Additional File 1, Suppl. Figure 2)**. To delve deeper into this relationship, we analyzed the TMB levels per chromatin modifier in comparison to canonical TMB markers such as the presence of DNA mismatch repair (MMR) mutations (*MLH1, MSH2, MSH6, PMS2*) and *POLE* or *POLD1* mutations. These results demonstrated higher TMB levels in cancers harboring alterations of certain chromatin modifiers (e.g. *DNMT3B, SMARCD1* and other) in comparison to cancers harboring mutations in canonical oncogenes/ tumor-suppressors, the MMR machinery or *POLE/POLD1* **(Figure 3A; Additional File 1, Suppl. Figure 18A)**. Next, we wanted to ascertain if the co-occurrence of mutations in the MMR machinery or *POLE/POLD1* together with alterations in chromatin modifiers would have an effect on TMB. Indeed, the co-occurrence of mutations in the MMR machinery or *POLE/POLD1* together with alterations in selected (high TMB CMs, medium TMB CMs, and low TMB CMs) chromatin modifiers had a cumulative effect, and the highest level of TMB **(Figure 3B; P<0.0001, Dunn’s test)**. To further refine this observation, we selected datasets harboring only MMR or *POLE/POLD1* mutations respectively, mutations only in high TMB CMs or combination of both. These analyses validated our previous observation that the presence of mutations in both chromatin modifiers and the MMR or *POLE/POLD1* machinery leads to a cumulative effect and the highest level of TMB **(Figure 3C and D, P<0.0001, Dunn’s test)**. Since high TMB levels have been associated with better response to immunotherapy with checkpoint inhibitors [14, 15], to assess the clinical relevance of our observations, we reanalyzed the data of non-small lung cell carcinoma patients treated with checkpoint inhibitors from the MSKCC cohort [16]. Patients harboring mutations in chromatin modifiers, which are associated with high TMB **(Additional File 1, Suppl. Figure 18A)** had a better durable effect compared to controls of all MSK genes or oncogenes/tumor-suppressors (P value, 0.028, and 0.029, respectively, Fisher’s exact test) **(Figure 3E; Additional File 1, Suppl. Figure 18B)**.

**Figure 3.**
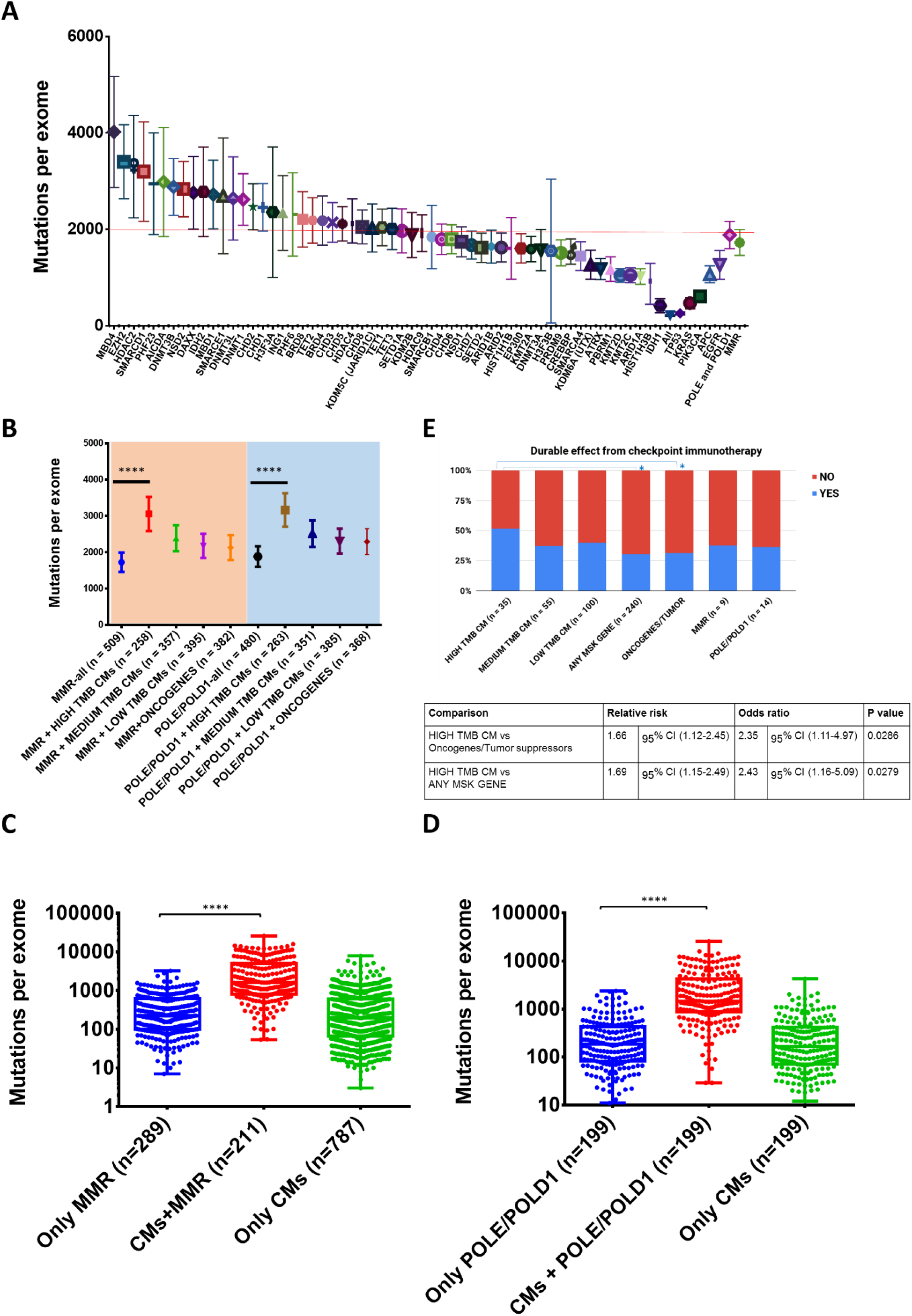
Tumor mutational burden analysis of TCGA samples. A) All samples harboring mutations in chromatin modifiers vs known oncogenes, tumor suppressors, MMR machinery and *POLE/POLD1*. B) All samples harboring mutations in the MMR machinery or *POLE/POLD1* (all samples) compared to a selection of MMR or *POLE/POLD1* samples accompanied by mutations in chromatin modifiers (arbitrarily divided in three groups). The P values can be found in Additional File 3. C) All samples harboring mutations only in the MMR machinery, high TMB chromatin modifiers (first third) and MMR machinery together, or only high TMB chromatin modifiers without mutations in the MMR machinery. D) All samples harboring mutations only in *POLE/POLD1*, high TMB chromatin modifiers (first third) and *POLE/POLD1* together, or only high TMB chromatin modifiers without mutations in *POLE/POLD1.* Asterisks in C and D indicate statistical significance (Dunn’s multiple comparisons test). E) A durable effect from checkpoint immunotherapy according to RECIST version 1.1 in 240 patients with NSCLC [16]. The cohort is subdivided into 7 groups: patients that harbor mutations in chromatin modifiers leading to high TMB (*SMARCD1-DAXX*), medium TMB (*KMT2A-H3F3B*), low TMB (*SMARCA4-IDH2*), and controls such as mutations in all MSK genes, mutations in selected oncogenes/tumor suppressors, mutations in the MMR machinery and mutations in POLE/POLD1. Asterisks indicate statistical significance (two-sided, Fisher’s exact test).

The presence of genomic alterations in chromatin modifiers have an effect on survival Finally, we wanted to evaluate the effect on the survival of patients harboring alterations in a chromatin modifier. We performed logistic regression analysis on TCGA data, and noticed that patients with mutations in the genes *SMARCD1, DNMT3L, TET3*, and *KMT2A* and amplifications in *EP300, KMT2B, KDM5C, DNMT3L, HDAC4, SMARCA4, ARID2* and *HDAC9* led to a worse survival prognosis, while mutations in *NSD1, CHD6, ARID1A, IDH1, IDH2* and amplifications in *HDAC2, IDH1* had a better survival prognosis **(Additional File 1, Suppl. Figure 19; Additional File 5)**.

## DISCUSSION

In this report, we sought to comprehensively study the features of thousands of cancer samples harboring genomic alterations in selected chromatin modifiers. Our study shows in detail that many of the pervasive genetic defects in cancer occur in the epigenetic machinery. Furthermore, the difference in the ubiquity of altered chromatin modifiers in some cancer types compared to others underlines the importance of epigenetic dysregulation for cancer formation and cancer progression. This means that some cancers might be more dependent on genetically induced epigenetic changes than others. From a mechanistic point of view, we discovered that in general, the presence of genomic alteration in one chromatin modifier is mutually exclusive with a genomic alteration in another. This observation illustrates that the underlying epigenetic landscape of cancer is highly selective and non-random, and alterations in too many essential chromatin factors might be unfavorable for the process of tumorigenesis. On the other hand, there are conspicuous cases where genetic aberrations in two or more chromatin modifiers co-occur together - which might point out to interesting interplays between major epigenetic networks (e.g. co-occurring amplifications of *DNMT1, SMARCA4*, and *BRD4* or co-occurring deletions in *HDAC2* and *ARID1B*). Approaches based on targeted dysregulation (genomic alteration) of one or multiple chromatin regulators coupled with functional analyses of the epigenome, might serve as an interesting basis for follow up studies, and novel targeted therapies. High tumor mutational burden (TMB) is an emerging biomarker of sensitivity to immune checkpoint inhibitors (PD-1 and PD-L1 blockade); Alterations in the DNA mismatch repair pathway and DNA polymerase ε have been used as a proxy for high TMB. Herein we showed that cancers harboring alteration in selected (not all) chromatin modifiers exhibit high TMB independently or MMR or *POLE/ POLD1*. In addition to that, we discovered that mutations in selected chromatin modifiers are associated with a more durable effect with checkpoint immunotherapy in patients with lung cancer. This find is quite relevant in the current developments of checkpoint immunotherapy and warrants further retrospective and prospective clinical studies. Finally, alterations in selected chromatin modifiers led to a worse/better patient prognosis, which in turn might encourage their development as prognostic markers in the future. All things considered, this study unveiled potential relationships between chromatin factors and cancer features and set the stage for additional follow up functional and clinical studies.

## METHODS

All genomic datasets reported in this study were downloaded from the cBioportal platform, http://www.cbioportal.org/, and statistically analyzed, and visualized in Microsoft Excel, Google Sheets, StatsDirect and Prism or other tools integrated within the cBioportal platform^11,12^. Supplementary Figures 2 and 3 were carried out on cancer datasets with >100 samples. The public cBioPortal is currently using hg19/GRCh37 version of the human reference genome. The main focus was put into MSK-IMPACT and TCGA datasets ^10,17^ https://www.cancer.gov/tcga.

Statistical analyses were performed using the Kruskal-Wallis exact test for multiple-group comparisons. Fisher’s exact test was used to compare outcomes. To compare survival rates, we applied the Kaplan-Meier estimator, and curves were compared by using the log-rank test. Odds ratios (OR) were calculated by logistic regression. Rank correlations were carried out by calculating the Spearman rank correlation coefficients.

## List of abbreviations

*CNA*: Copy Number Amplification
*TCGA*: The Cancer Genome Atlas
*MSK-IMPACT*: Memorial Sloan Kettering Integrated Mutation Profiling of Actionable Cancer Targets
*MMR*: DNA Mismatch Repair
*PD-1*: Programmed cell death protein 1
*PD-L1*: Programmed death-ligand 1

## Declarations

### Ethics approval and consent to participate

All data are taken from publicly available datasets and cancer projects.

### Consent for publication

Not applicable. All data are taken from publicly available datasets and already reported in the literature.

### Availability of data and material

All data are taken from the cBioportal, http://www.cbioportal.org.

### Authors’ contributions

G.K conceived and designed the study. G.K and M.M processed, analyzed and interpreted the data. R.R helped in data analysis and statistics. A.S, S.M, M.J, Z.M, and I.K helped in data analysis and interpretation. G.K wrote the manuscript. All authors contributed to the improvement of the manuscript and read the final version of the manuscript.

## Acknowledgments

Not applicable.

