## Additional File 1 for "Genetic aberrations in chromatin factors are pervasive, mutually exclusive, associated with a high mutational burden and improved response to checkpoint immunotherapy in cancer"

**Corresponding author*

**A**

| Chromatin remodelers |
|---|
| *SMARCB1* |
| *SMARCA4* |
| *PBRM1* |
| *ARID1A* |
| *ARID1B* |
| *ARID2* |
| *SMARCD1* |
| *SMARCE1* |
| *ATRX* |
| *DAXX* |
| *CHD5* |
| *CHD2* |
| *CHD1* |
| *CHD3* |
| *CHD4* |
| *CHD6* |
| *CHD7* |
| *CHD8* |

| Histone de/methylation |
|---|
| *KMT2A* |
| *KMT2D* |
| *KMT2C* |
| *SETD1A* |
| *PRDM9* |
| *EZH2* |
| *NSD1* |
| *NSD2* |
| *SETD2* |
| *KDM5C (JARID1C)* |
| *KDM6A (UTX)* |
| *KDM2B* |

| Histones |
|---|
| *H3F3A* |
| *H3F3B* |
| *HIST1H3B* |
| *HIST1H1B* |

| Histone de/acetylation |
|---|
| *EP300* |
| *CREBBP* |
| *HDAC2* |
| *HDAC4* |
| *HDAC9* |

| Readers |
|---|
| *PHF6* |
| *PHF23* |
| *BRD4* |
| *BRD8* |
| *ING1* |

| DNA de/methylation | |
|---|---|
| *DNMT3A* | *TET2* |
| *DNMT3B* | *TET1* |
| *DNMT3L* | *TET3* |
| *DNMT1* | *IDH1* |
| *AICDA* | *IDH2* |
| *MBD1* | |
| *MBD4* | |

**B**

| Chromatin remodeling |
|---|
| *SMARCB1* |
| *SMARCA4* |
| *PBRM1* |
| *ARID1A* |
| *ARID1B* |
| *ARID2* |
| *SMARCD1* |
| *ATRX* |
| *DAXX* |

| DNA de/methylation |
|---|
| *DNMT3A* |
| *DNMT3B* |
| *DNMT1* |
| *TET2* |
| *TET1* |
| *IDH1* |
| *IDH2* |

| Histone methylation/ acetylation |
|---|
| *EP300* |
| *CREBBP* |
| *KMT2A* |
| *KMT2D* |
| *KMT2C* |
| *EZH2* |
| *NSD1* |
| *SETD2* |
| *KDM5C* |
| *KDM6A* |

| Histone proteins |
|---|
| *H3F3A* |
| *H3F3B* |
| *HIST1H3B* |

| Histone PTM reader |
|---|
| *BRD4* |

**Supplementary Figure 1**. List of chromatin modifiers evaluated in this study. A) Non-redundant datasets deposited in the cBioportal including TCGA datasets B) MSK-IMPACT study.

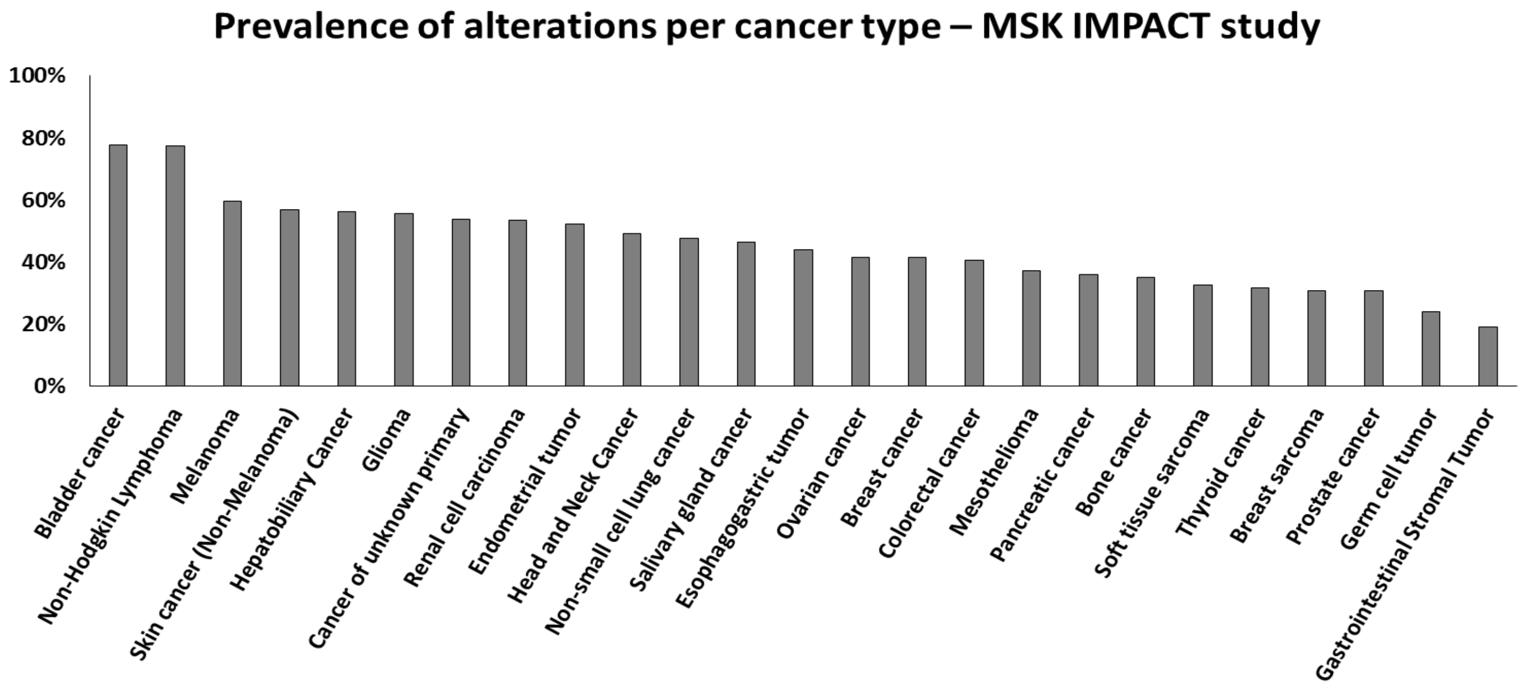


**Supplementary Figure 2**. Prevalence of alterations in chromatin modifiers per cancer type in the MSK-IMPACT study.

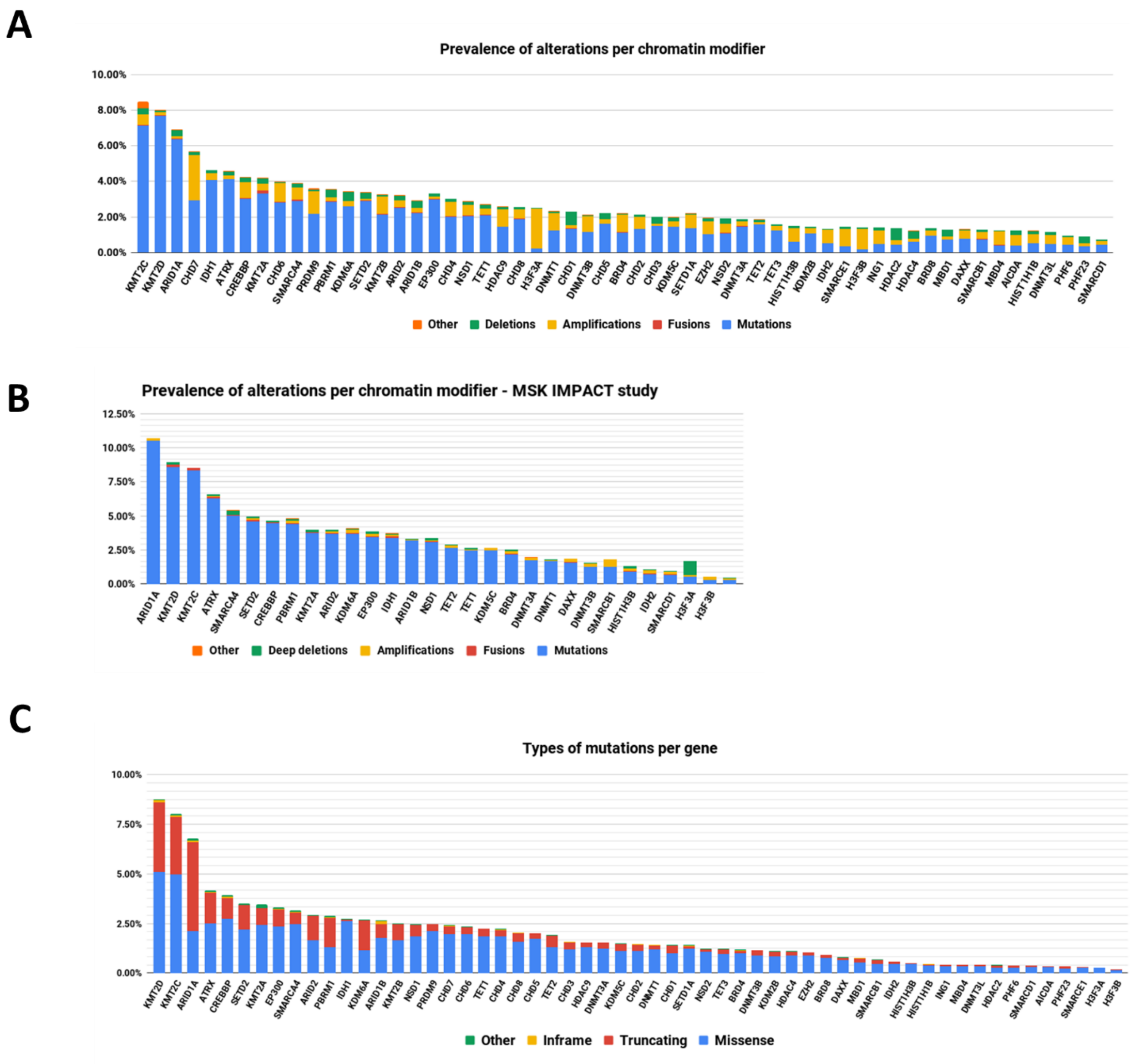


**Supplementary Figure 3**. Types of genomic alterations per chromatin modifier. A) Non-redundant datasets deposited in the cBioportal including TCGA datasets B) MSK-IMPACT study C) Types of mutations in non-redundant studies.

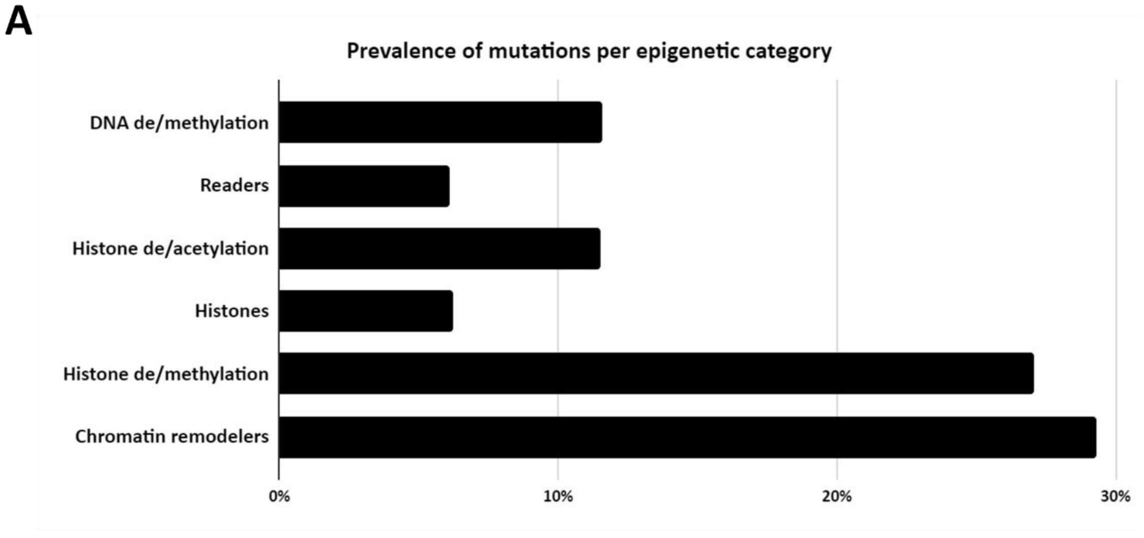


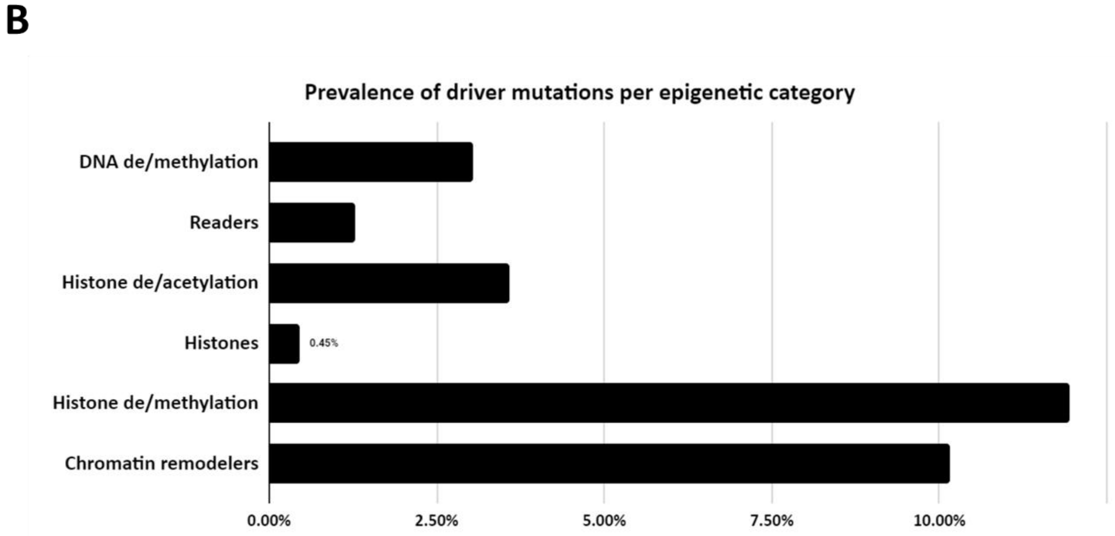


**Supplementary Figure 4**. Frequency of genomic alterations per epigenetic category enlisted in Supplementary Figure 1. A) Putative driver + driver + VUS alterations B) Putative driver + driver alterations.

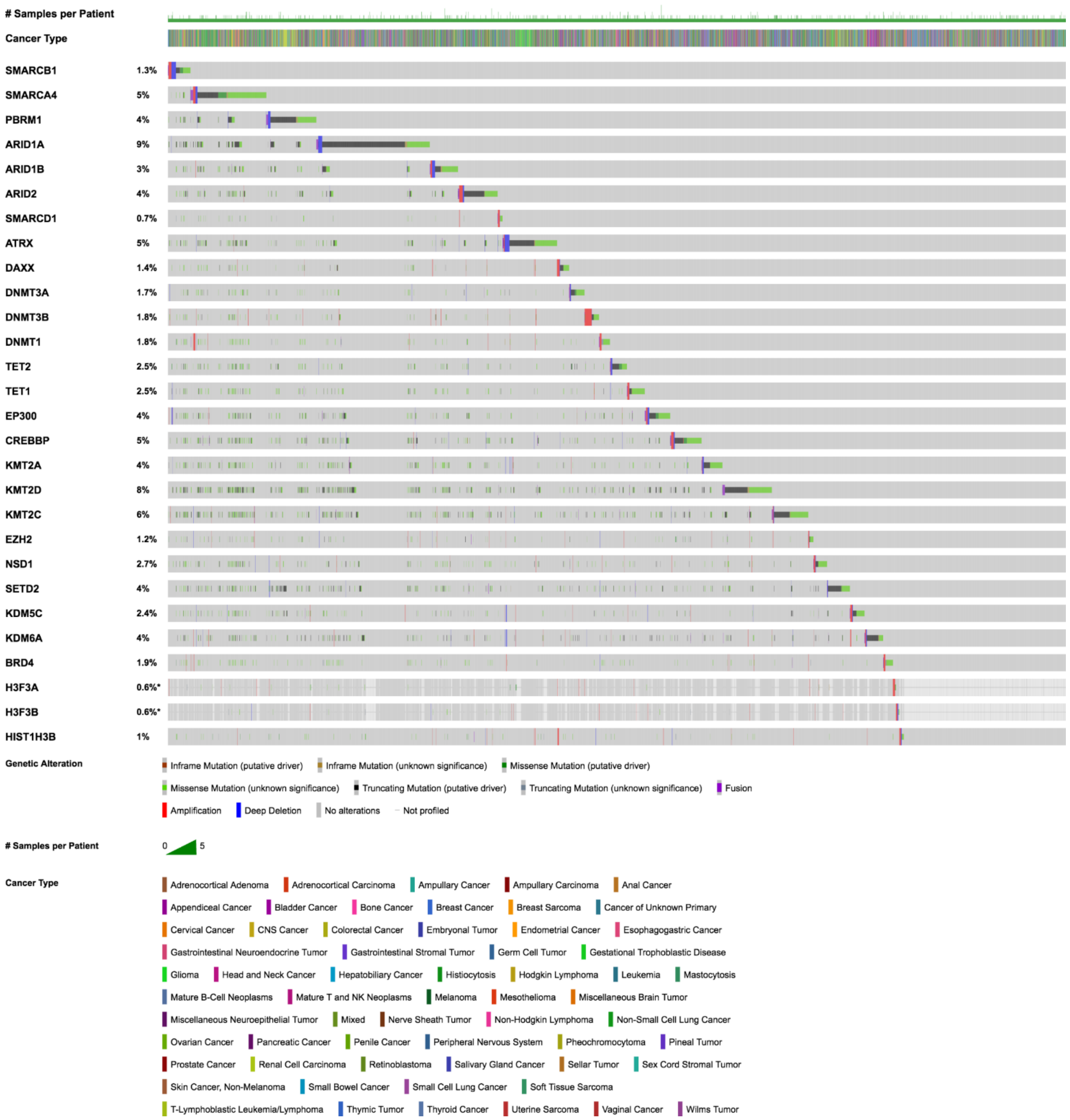


**Supplementary Figure 5.** Oncoprint analysis of alterations in chromatin modifiers in samples from the MSK-IMPACT study.

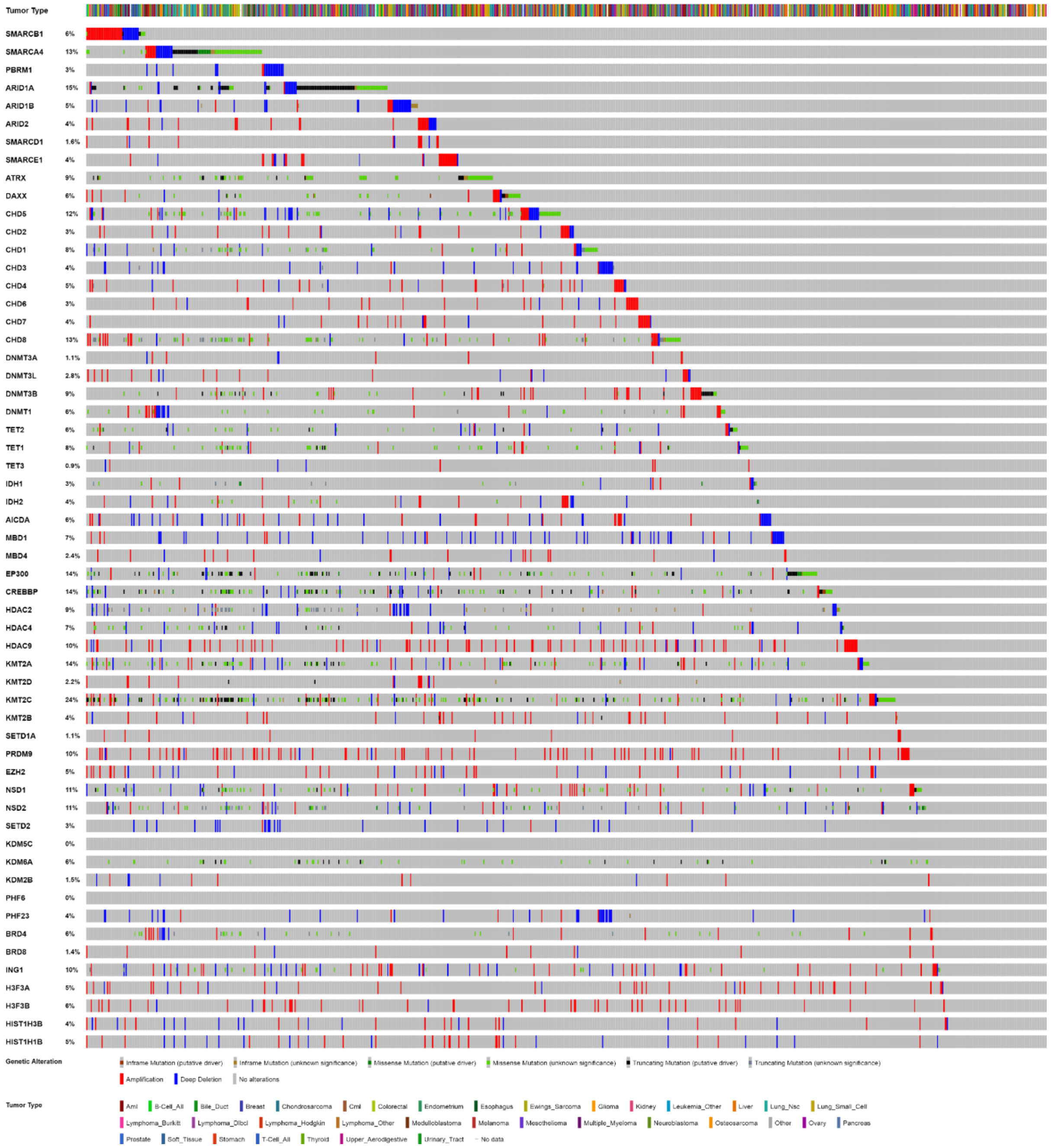

**Supplementary Figure 6**. Oncoprint analysis of alterations in chromatin modifiers in cancer cell lines from Cancer Cell Line Encyclopedia.

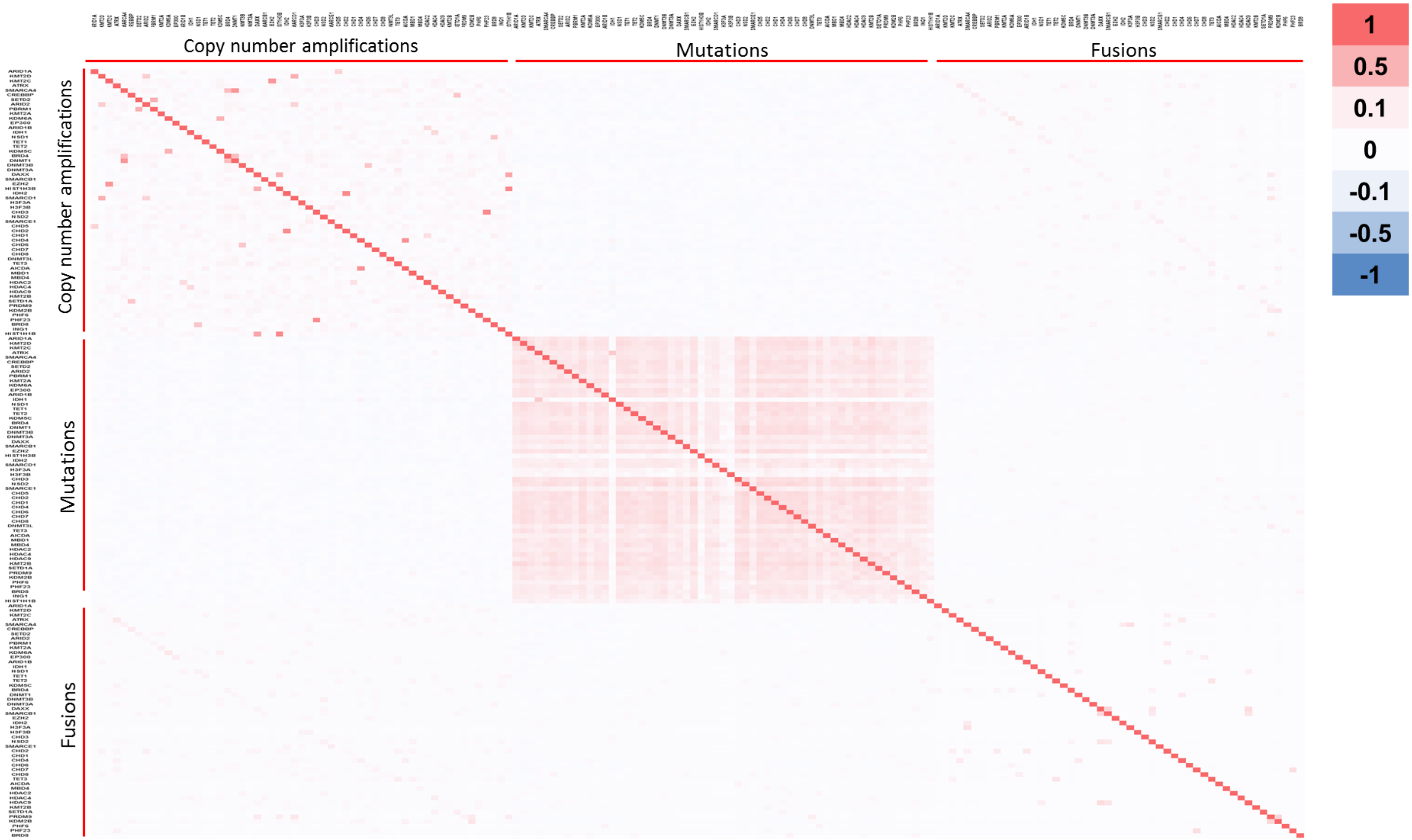


**Supplementary Figure 7**. Spearman correlation coefficient analysis of alterations in chromatin modifiers taken from the TCGA study. More details in Additional File 2.

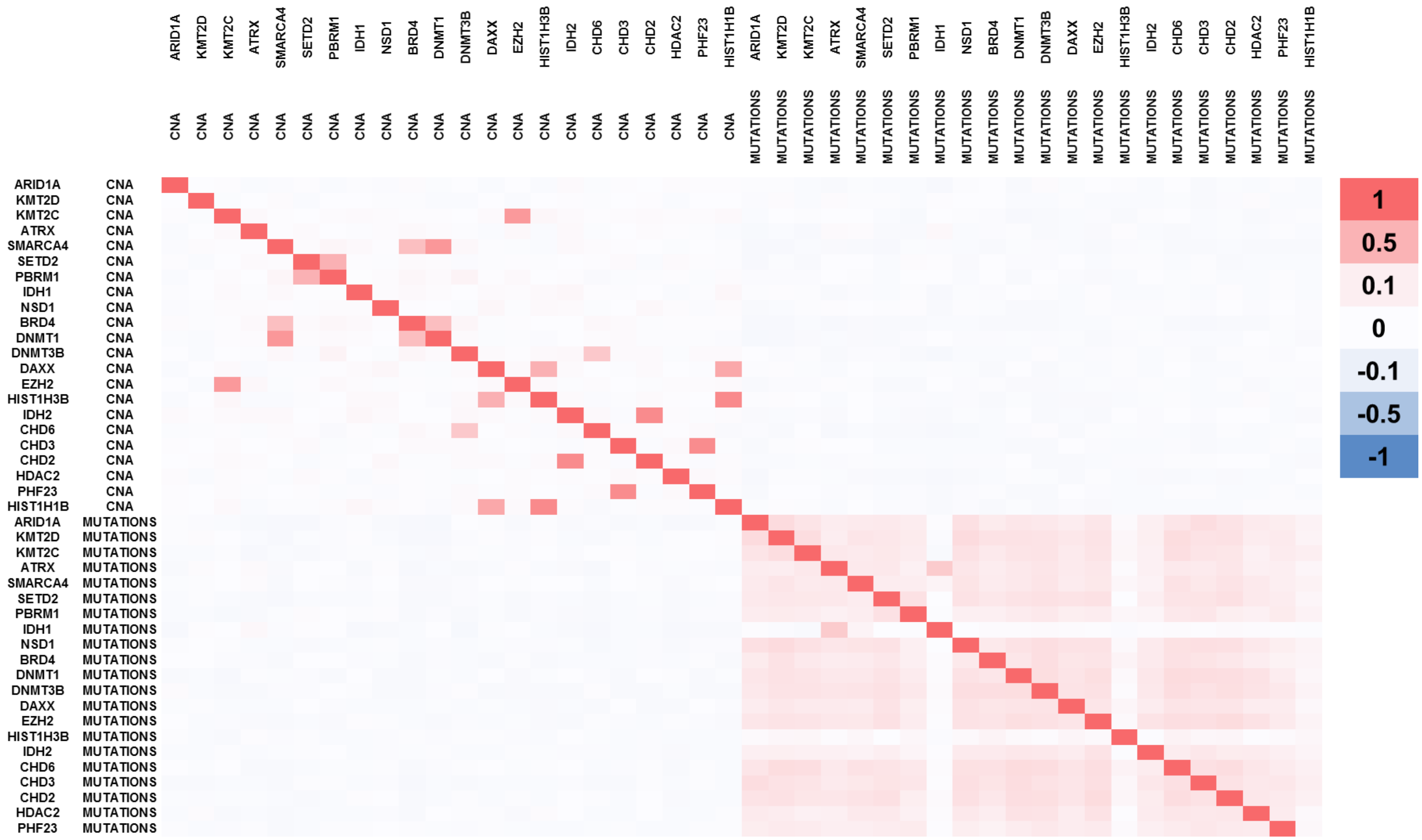


**Supplementary Figure 8**. Spearman correlation coefficient analysis of alterations in selected chromatin modifiers taken from the TCGA study. More details in Additional File 2.

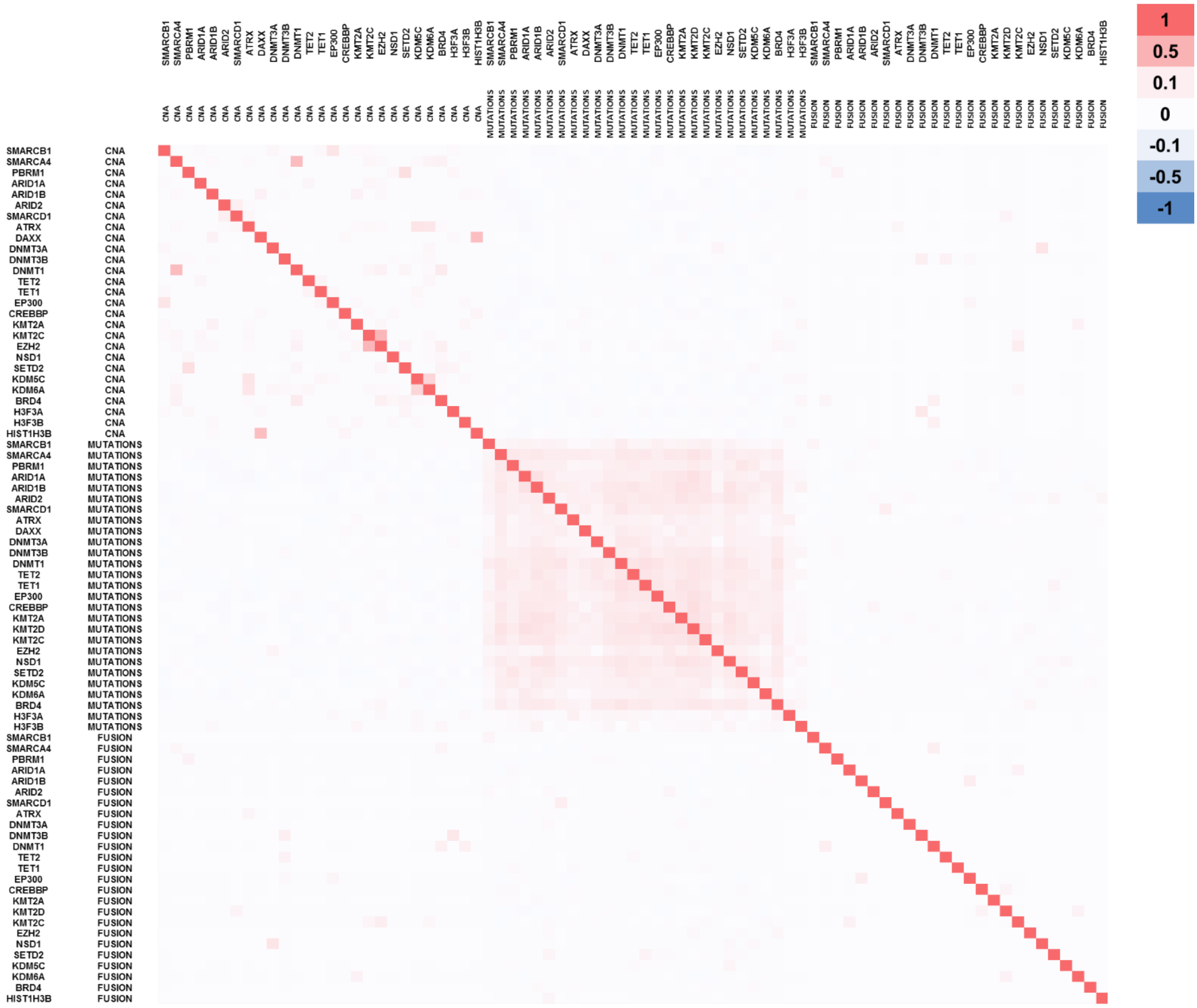


**Supplementary Figure 9**. Spearman correlation coefficient analysis of alterations in chromatin modifiers taken from the MSK-IMPACT study. More details in Additional File 2.

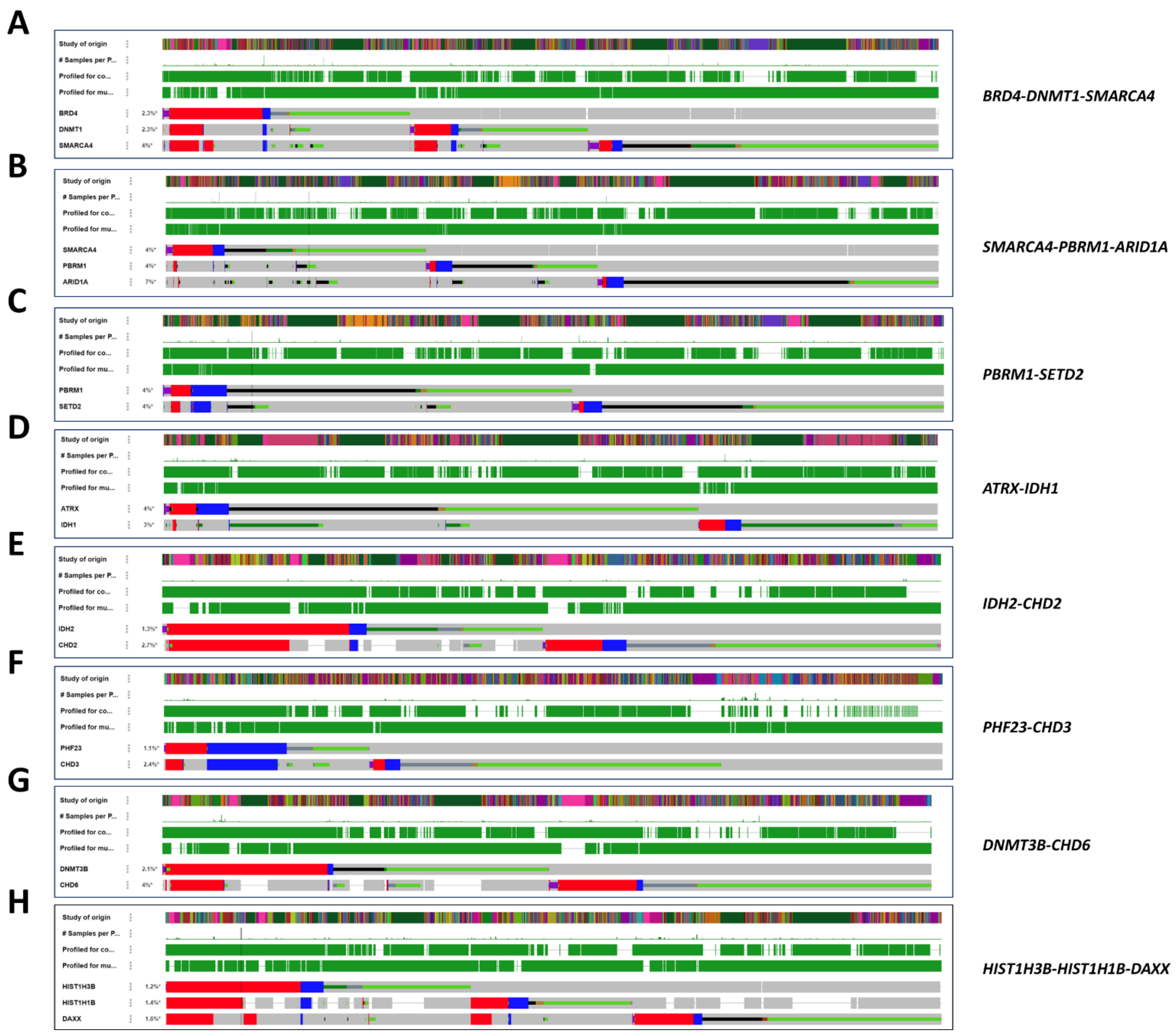


**Supplementary Figure 10**. Oncoprint analysis showing co-occurrence of alterations in chromatin modifiers in samples taken from the TCGA study. A) *BRD4-DNMT1-SMARCA4*, B) *SMARCA4-PBRM1-ARID1A*, C) *PBRM1-SETD2*, D) *ATRX-IDH1*, E) *IDH2-CHD2*, F) *PHF23-CHD3*, G) *DNMT3B-CHD6*, H) *HIST1H3B-HIST1H1B-DAXX.*

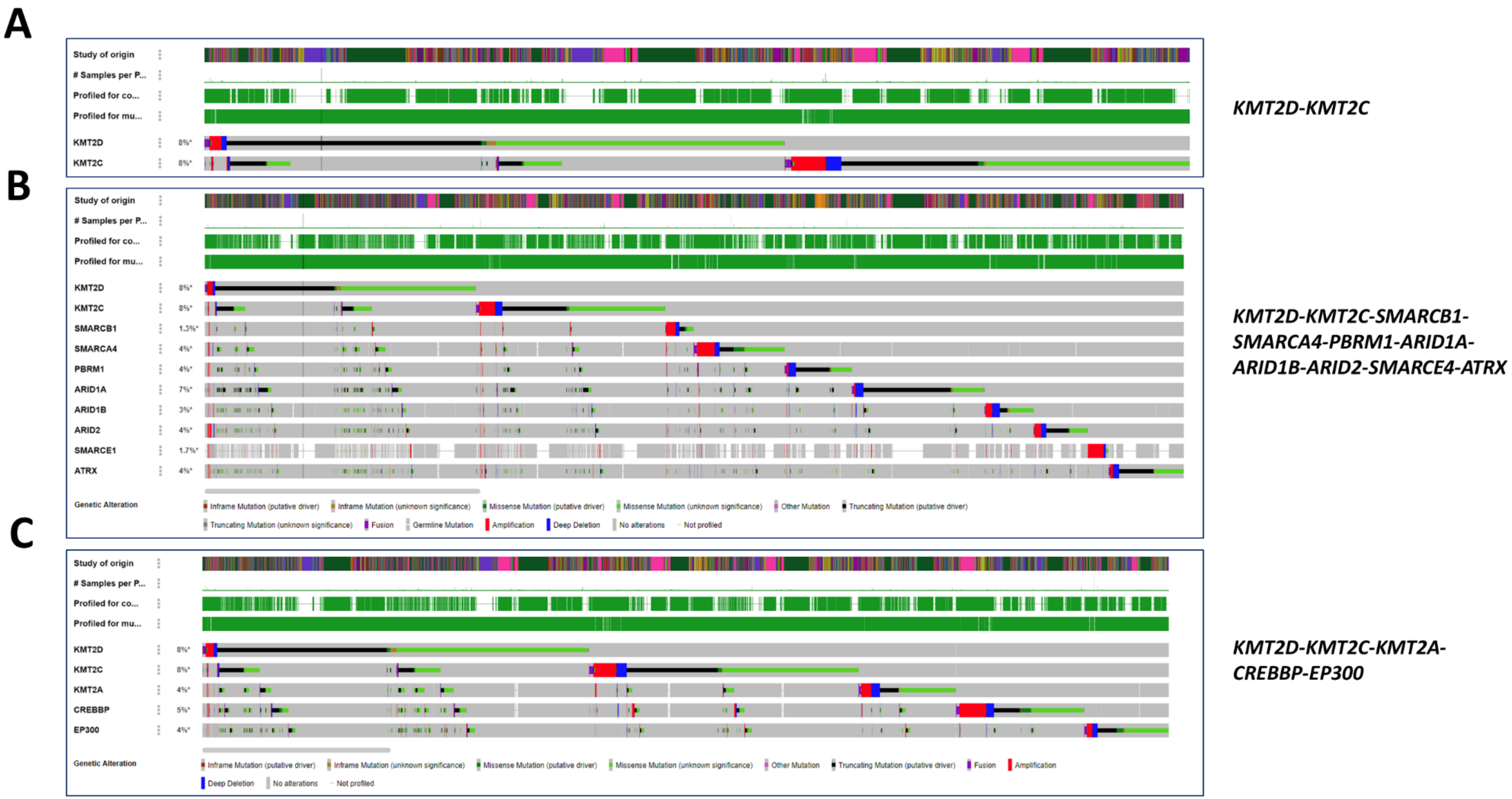


**Supplementary figure 11**. Oncoprint analysis showing co-occurrence of alterations in chromatin modifiers in samples taken from the TCGA study. A) *KMT2D-KMT2C*, B) *KMT2D-KMT2C-SMARCB1-SMARCA4-PBRM1-ARID1A-ARID1B-ARID2-SMARCE4-ATRX*, C) *KMT2D-KMT2C-KMT2A-CREBBP-EP300.*

#### Lung cancer

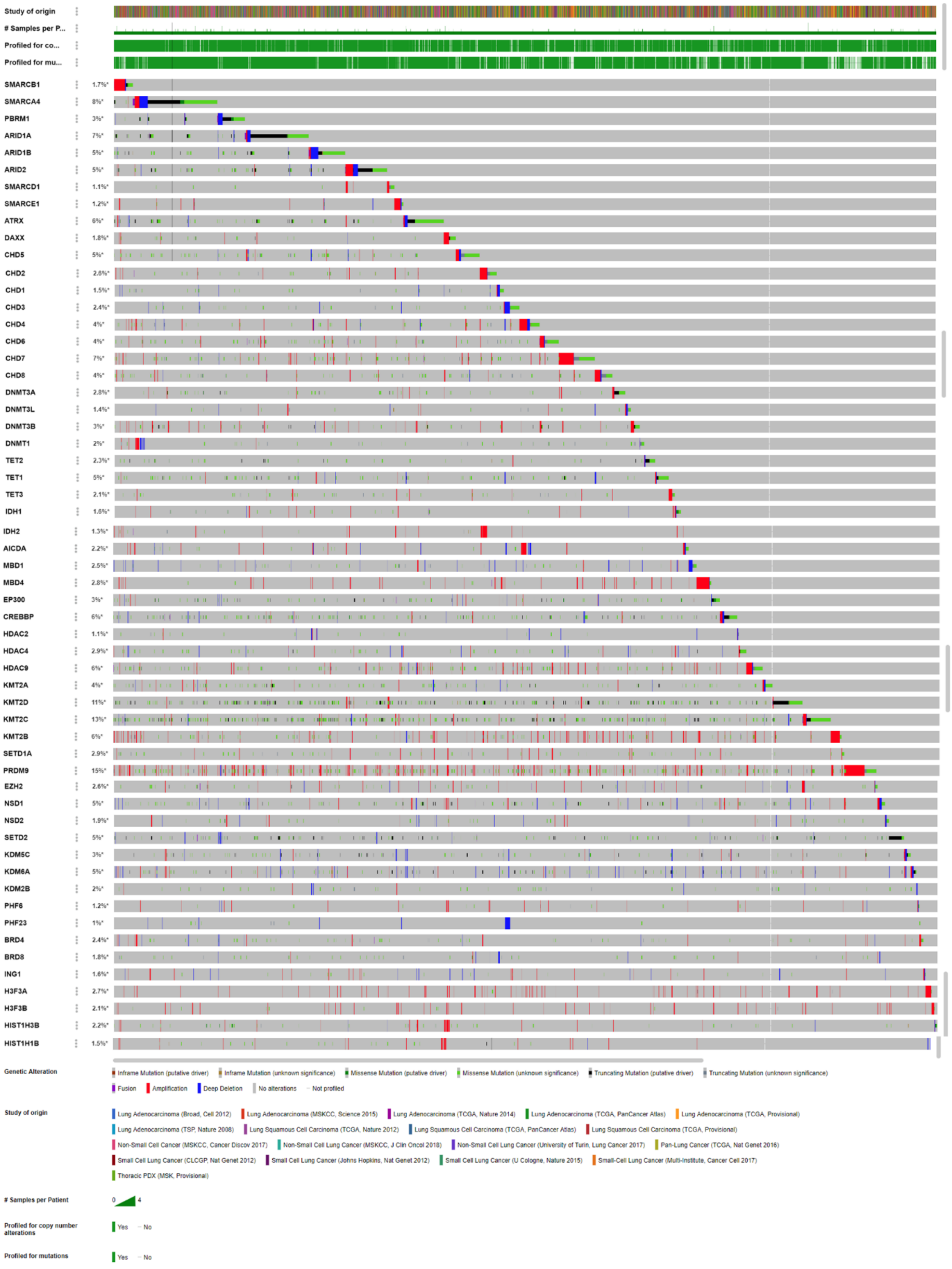


**Supplementary figure 12**. Oncoprint analysis of alterations in chromatin modifiers in samples of lung cancer.

#### Bladder cancer

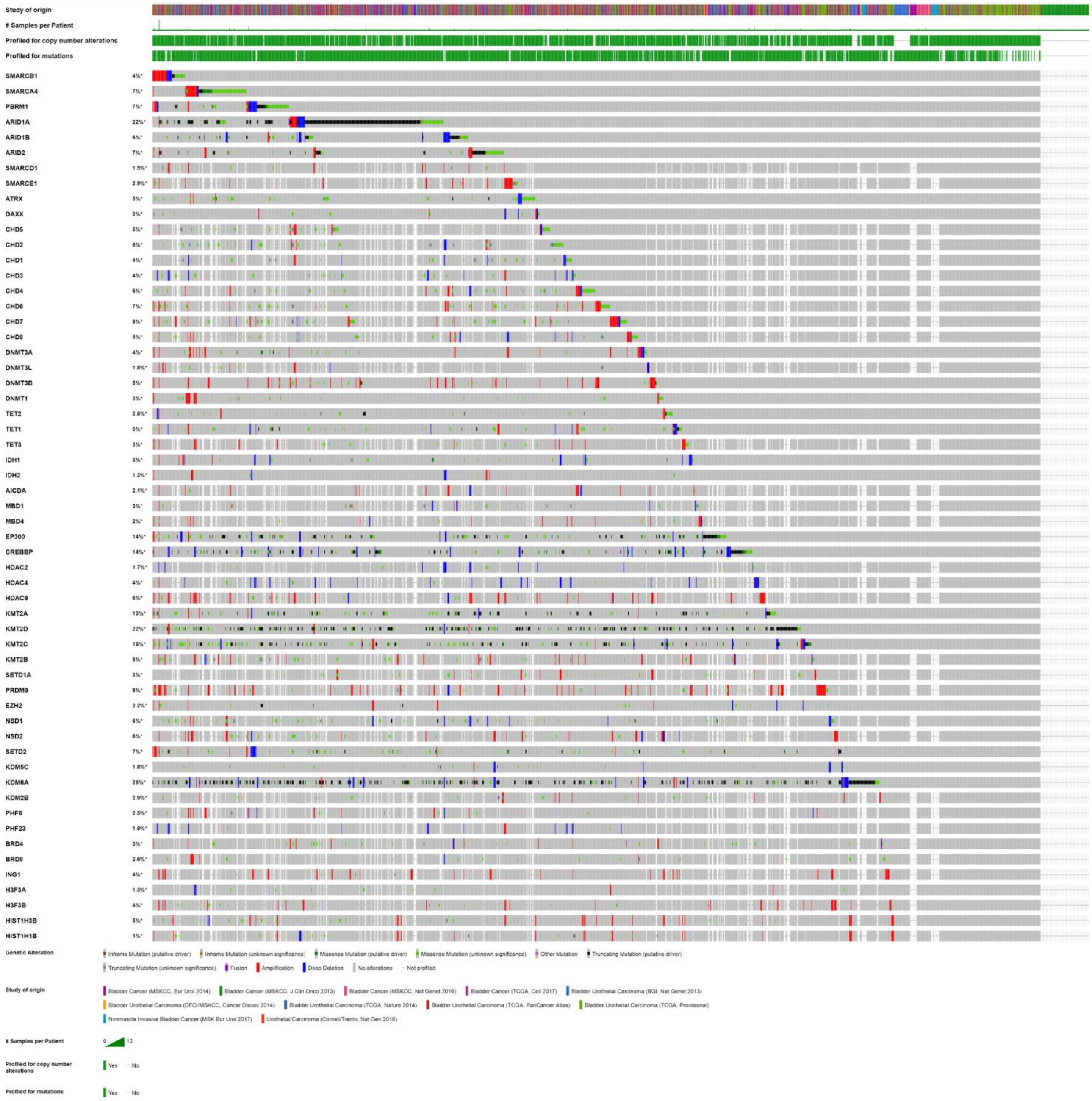


**Supplementary figure 13**. Oncoprint analysis of alterations in chromatin modifiers in samples of bladder cancer.

### Low grade glioma and glioblastoma

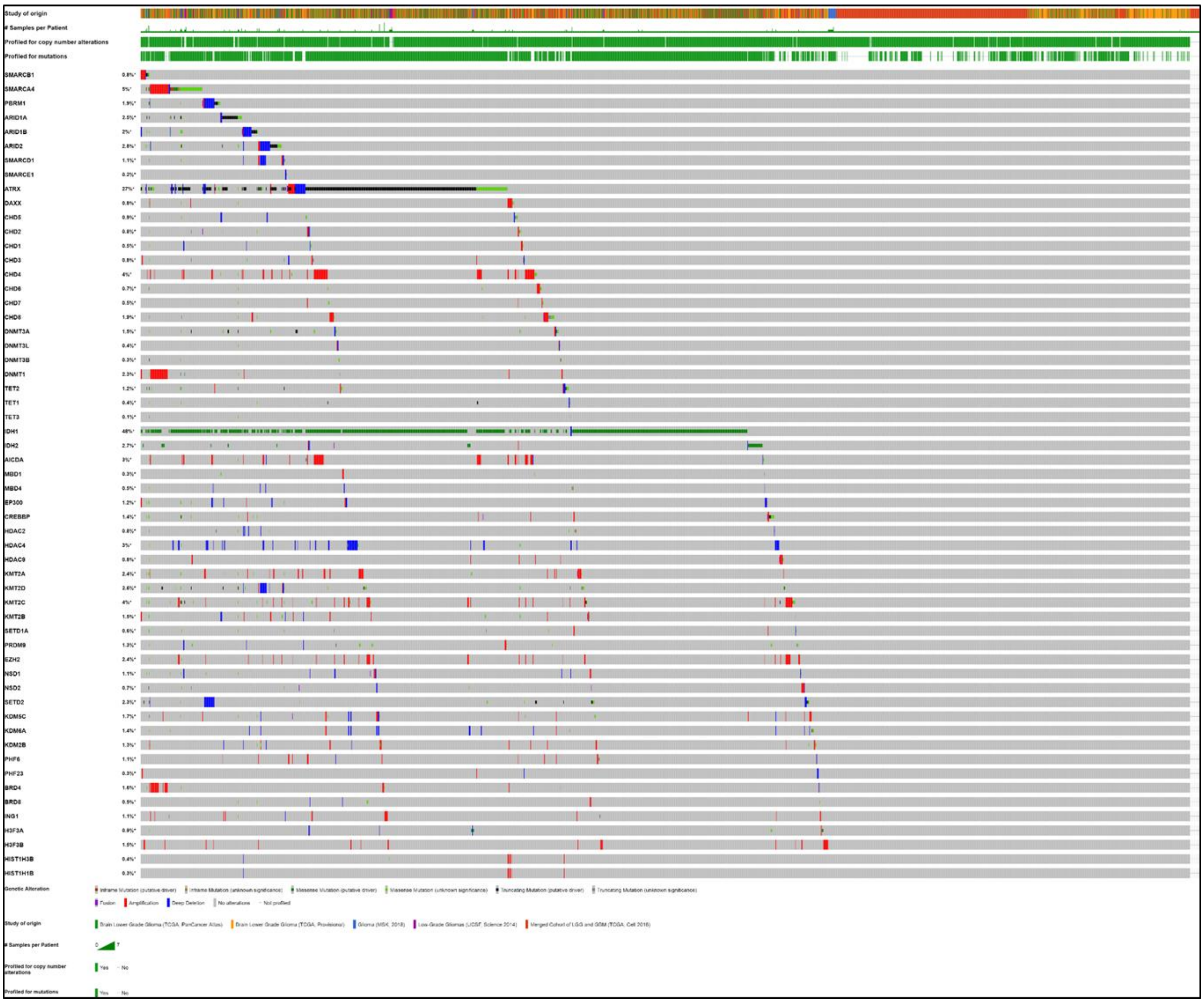


**Supplementary figure 14**. Oncoprint analysis of alterations in chromatin modifiers in samples of low-grade glioma and glioblastoma.

### Melanoma

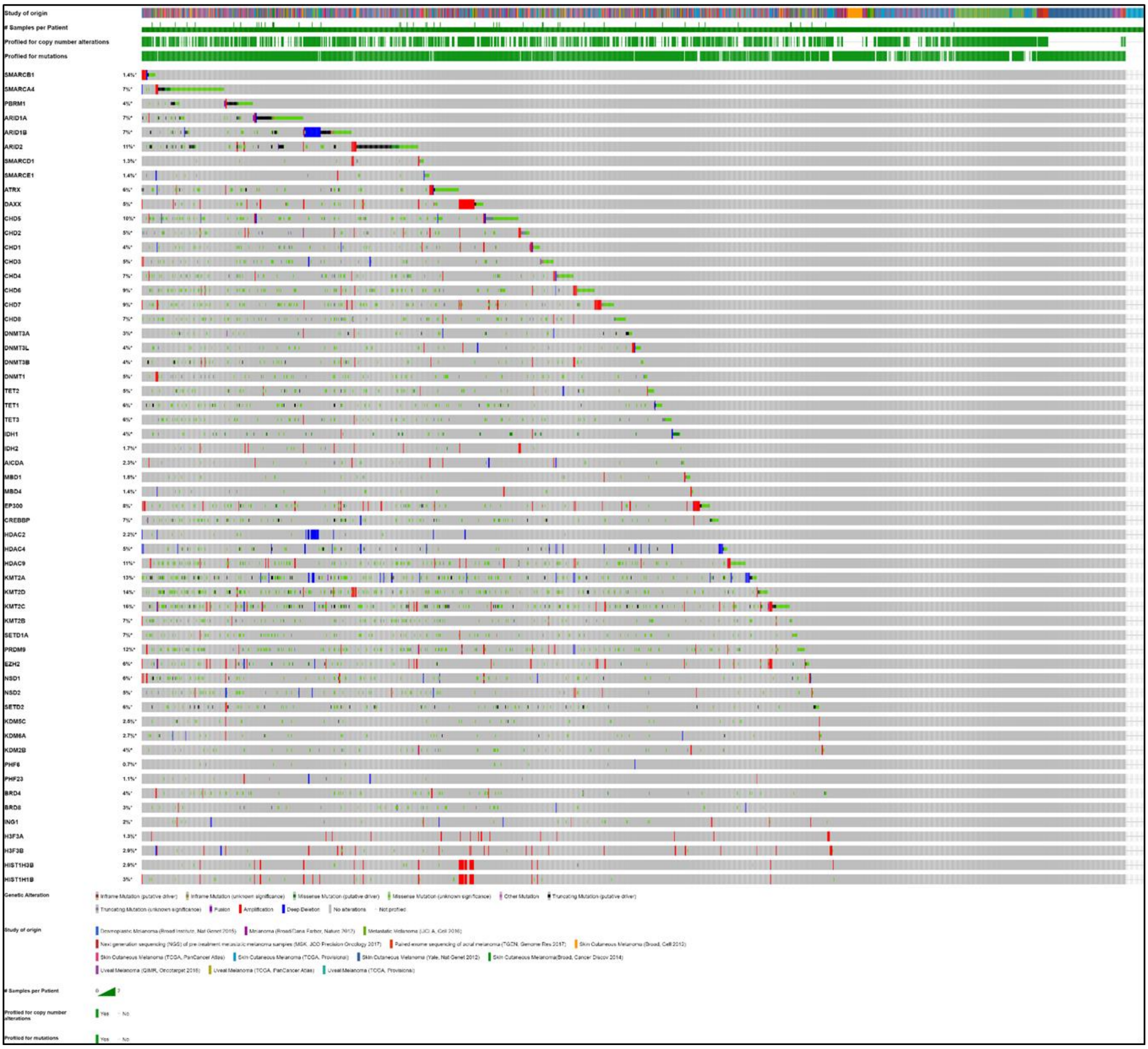


**Supplementary figure 15**. Oncoprint analysis of alterations in chromatin modifiers in samples of melanoma.

### Endometrial and cervical cancer

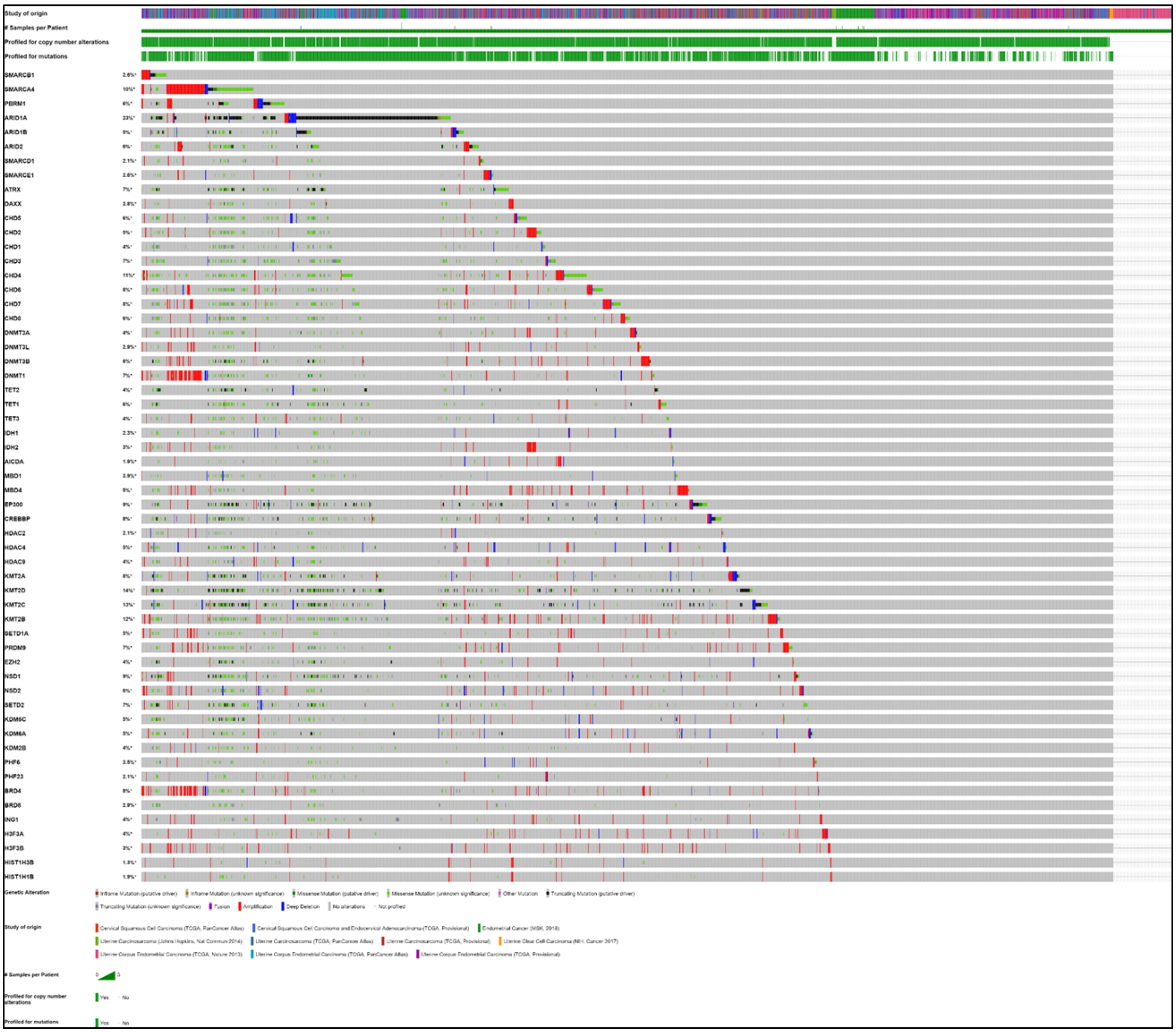

**Supplementary figure 16**. Oncoprint analysis of alterations in chromatin modifiers in samples of endometrial and cervical cancer.

A

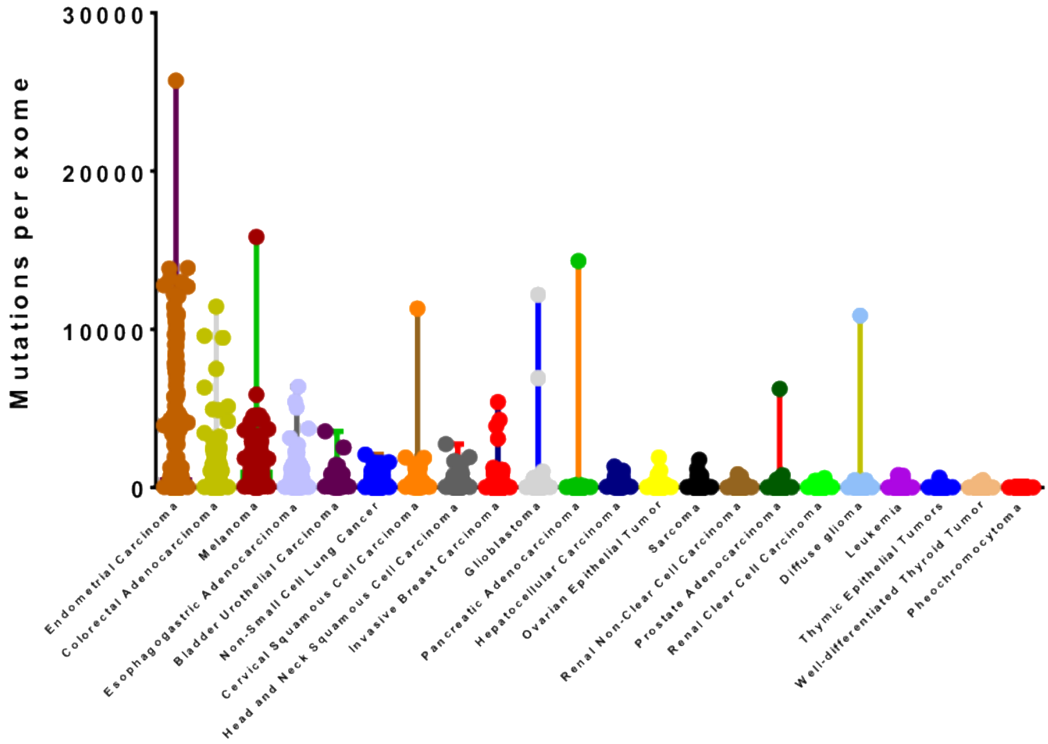


B

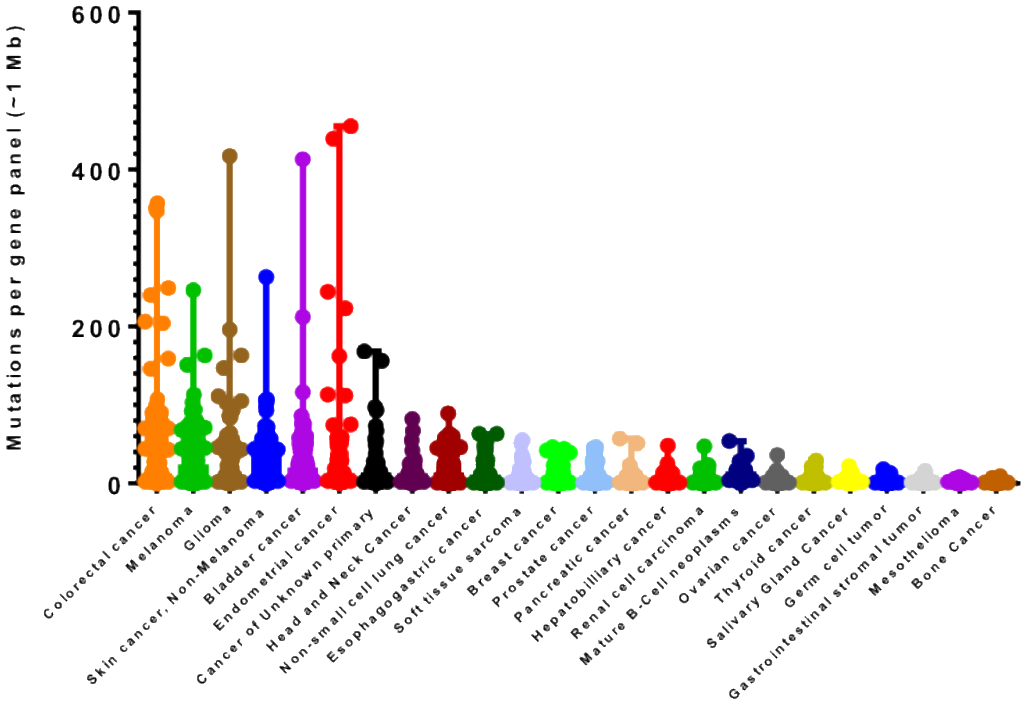


**Supplementary figure 17**. Analysis of tumor mutational burden (TMB). A) Maximal and minimal values of TMB per cancer type from the TCGA study B) Maximal and minimal values of TMB per cancer type from the MSK-IMPACT study.

**A**

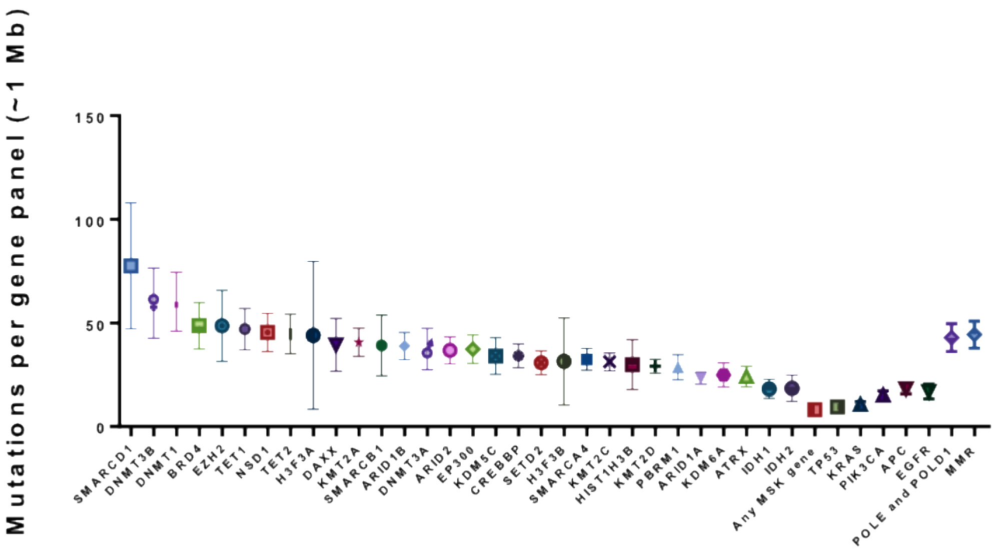


**B**

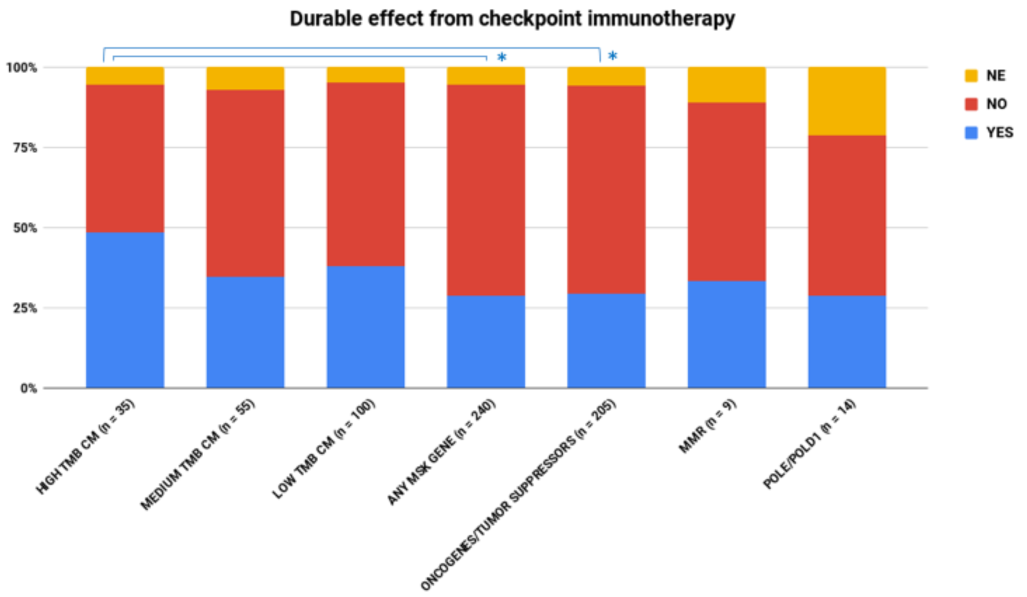


**Supplementary figure 18**. A) Shown is the mean and 95% CI of TMB per cancer samples harboring the alterations in chromatin modifiers, all MSK-IMPACT genes, common oncogenes, and tumor suppressors, *POLE/POLD1* genes, and MMR genes. The p values can be found in Additional File 3. B) A durable effect from checkpoint immunotherapy according to RECIST version 1.1 in 240 patients with NSCLC (Rizvi, 2018). The cohort is subdivided into 7 groups: patients that harbor mutations in chromatin modifiers leading to high TMB (SMARCD1-DAXX), medium TMB (KMT2A-H3F3B), low TMB (SMARCA4-IDH2), and controls such as mutations in all MSK genes, mutations in selected oncogenes/tumor suppressors, mutations in the MMR machinery and mutations in *POLE/POLD1*. Asterisks indicate statistical significance (two-sided, Fisher's exact test).

A

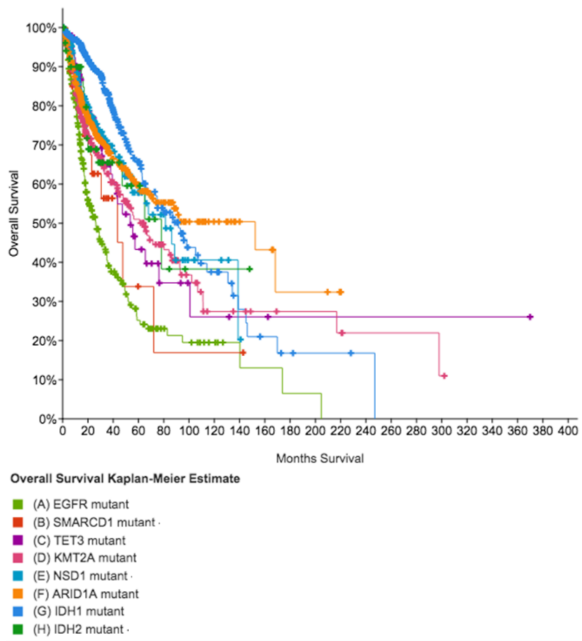


B

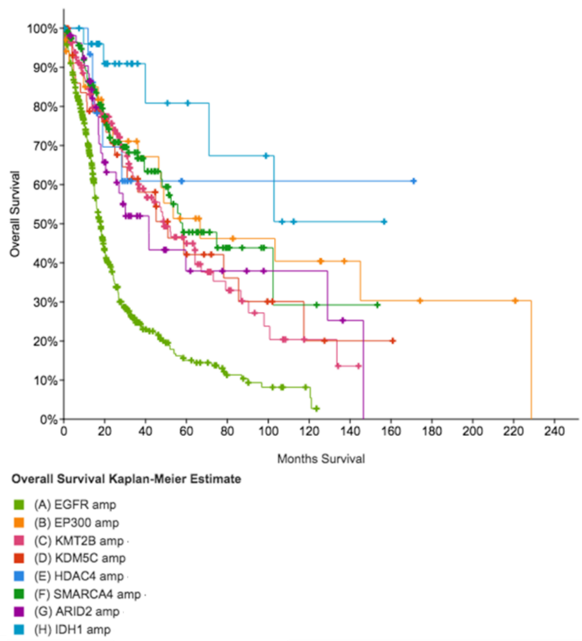


**Supplementary figure 19.** Kaplan-Meier graphs of altered chromatin modifiers leading to a worse/better prognosis selected from Additional File 5. A) Mutations in chromatin modifiers. *EGFR* is used as a representative oncogene for comparison B) Amplifications in chromatin modifiers. *EGFR* is used as a representative oncogene for comparison.
